# Multi-modal Graph Integration for Biologically Interpretable Domain Identification Using Spatial Transcriptomics

**DOI:** 10.64898/2026.07.30.741921

**Authors:** Seungeun Lee, Guolon Wang, Kyungtae Kang, Jichao Chen, Mingon Kang

## Abstract

Functional domain identification in spatial transciptomics transforms spatial molecular measurements into mechanistic insights into tissue physiology and pathology. However, the inherent noise and sparsity of gene expression data, along with the locality-biased design of conventional graph-based approaches, fundamentally limit the accurate identification of complex tissue domains. In this study, we propose a novel Biologically Interpretable multi-modal Graph using Spatial Transcriptomics, called BIGraph-ST, that integrates pathway activity scores and histological image features for robust spatial domain identification. BIGraph-ST represents modality-specific similarity through affinity graphs and propagates spatial topology to capture higher-order connectivity within the tissue microenvironment. Experimental results demonstrated robust performance and notable improvements across multiple gold-standard benchmark datasets, particularly in cancer tissues. Moreover, BIGraph-ST provides biologically interpretable pathway-level representations of domains, which ultimately offers a valuable tool to gain biological insights into complex tissue architectures. The source code will be publicly available upon acceptance.

## 1 Introduction

Domain identification using spatial transcriptomics is central to understanding tissue architecture and molecular programs that jointly define biological functions. Spatial transcriptomics integrates traditional histology with gene expression while preserving spatial organization, which allows computational delineation of such domains [11, 12]. Spatial transcriptomics-based domain identification not only aligns with established anatomy but also provides a principled reduction of the hypothesis space for downstream cell-level analyses, including cell typing and deconvolution. This capability enables systematic characterization of tissue heterogeneity and intercellular signaling, facilitating the discovery of disease biomarkers and therapeutic targets [7, 17]. Methodologically, domain-level analysis leverages spatial learning and neighborhood aggregation to improve robustness under pervasive gene sparsity, which is a fundamental challenge in high-resolution spatial transcriptomics.

Deep learning models have demonstrated substantial potential in unsupervised clustering tasks for spatial domain identification. In spatial transcriptomics, spatial context is typically modeled as a graph, where nodes represent tissue spots with node features of gene expression profiles, and edges encode spatial proximity. Graph neural networks operate on this graph structure and propagate genetic information across neighboring nodes to learn spatially-based representations. Current approaches for domain identification can be categorized into bifold: (i) contrastive learning and (ii) autoencoder-based embedding frameworks. Contrastive learning methods learn spot representations by putting embeddings of spatially similar spots together while pushing apart dissimilar spots. For instance, GraphST [10] applies corrupted graph augmentations as negative pair with a graph-level contrastive loss, and SpaGIC [9] iteratively refines domain-aware spot embeddings and leverages pseudo-labels to sharpen spatial boundaries. On the other hand, autoencoder-based frameworks learn low-dimensional embeddings of spatial transcriptomic spots using reconstruction losses. STAGATE [5] employs a graph attention autoencoder to aggregate information from spatial neighbors, and SEDR [16] utilizes a deep autoencoder coupled with a variational graph autoencoder to embed spatial information into gene expression representations.

Recent studies have explored integrating histological images as complementary morphological information into spatial transcriptomic analysis [2]. SpaGCN refines spatial edge weights using morphological similarity through correlations of RGB intensities between spots [7]. DeepST augments gene expression with deep learning based image features in node attributes [15]. ConST aligns modalities using cross-modal contrastive learning between transcriptomic and visual embeddings [19]. Despite these advances, such approaches incorporate histology as auxiliary to spatial graphs or node features, without explicitly modeling inter- and intra-modality relationships. Furthermore, most current approaches rely on locally defined spatial graphs that connect immediate neighbors rather than capture global tissue organization. This locality bias may limit their ability to represent complex connectivity in tissue microenvironment [3].

A further challenge arises from the raw gene expression profiles in spatial transcriptomics. The inherent sparsity and noise in gene expression profiles make genomic representations vulnerable to instability and spurious signals [13, 14]. Pathway scoring provides an alternative representation of gene expression by grouping genes into functionally associated sets, which captures hierarchical interactions while reducing the impact of sparsity and spot-level noise. For instance, PathMGCN constructs a functional similarity graph using pathway activity scores, while jointly learning a spatial proximity graph [18]. However, the latent representations of PathMGCN are primarily driven by gene-level reconstruction, which consequently limits the effective use of pathway-level semantics. Additionally, PathMGCN relies solely on transcriptomic data without morphological features.

In this study, we propose a multimodal graph diffusion framework, named BIGraph-ST, for functionally interpretable spatial domain identification. Unlike existing approaches that are mainly based on spatial proximity to construct graphs, BIGraph-ST represents functional and morphological relationships by constructing modality-specific graphs and diffusing node attributes to capture higher-order tissue organization. This joint modeling of attribute-level similarities and spatial topology provides robust domain identification that aligns with both pathways and tissue morphology. Our contributions are summarized as follows: (i) BIGraph-ST integrates comprehensive connectivity driven by pathways and histology, which has been overlooked by conventional spatial proximity–based methods, (ii) BIGraph-ST can capture higher-order tissue connectivity, overcoming the locality bias of the current approaches, and (iii) Pathway-level representations in BIGraph-ST enable biologically meaningful functional characterization in domain identification.

## 2 Method

BIGraph-ST integrates each spot’s pathway activity, histological image features, and spatial context within a tissue to identify functional spatial domains. BIGraph-ST consists of four key steps: (i) construction of modality-specific spectral embeddings, (ii) spatial topology-aware diffusion, and (iii) cross-modal affinity integration, and (iv) domain clustering (Fig. 1). First, BIGraph-ST constructs modality-specific spectral embeddings (i.e., pathway- and histology-derived affinity graphs) to capture attribute-level similarity among spots. Second, spatial topology-aware diffusion propagates these spectral embeddings over the spatial graph to model long-range spatial dependencies. Third, cross-modal integration refines the spatially smoothed embeddings through multi-modal affinity graph diffusion. Finally, BIGraph-ST identifies functional spatial domains using *k*-means clustering.

**Fig. 1:**
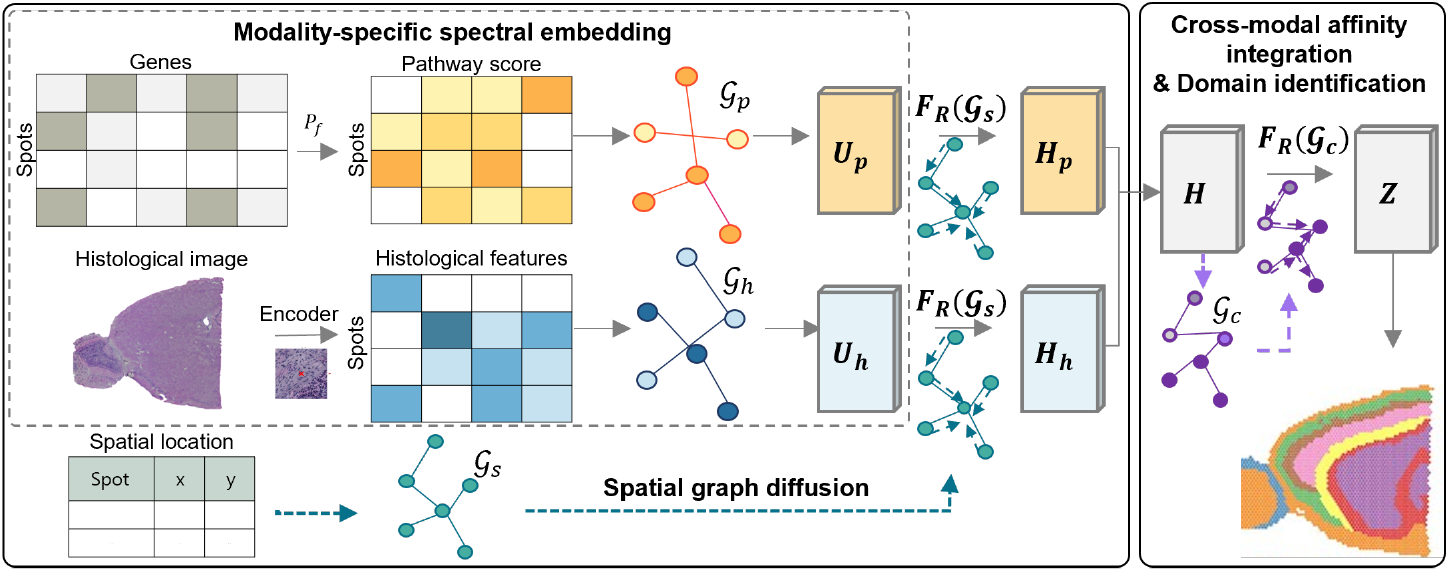
Overview of BIGraph-ST for identifying functional spatial domains. The framework consists of four steps: (i) extracting modality-specific spectral embeddings to capture attribute-level similarities; (ii) applying spatial graph diffusion to model long-range spatial dependencies; (iii) refining embeddings using crossmodal diffusion, and (iv) applying k-means clustering to identify meaningful spatial domains.

Spatial transcriptomics data comprise a gene expression matrix **G**, a histological image **I**, and spatial coordinates of spots **C**. Specifically, **G** ∈ ℝ^*n×g*^ denotes the gene expression matrix, and 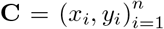 represents the spatial location of each spot *i*. Here, *n* and *g* denote the number of spots and genes, respectively, and each row **g**_*i*_ ∈ ℝ^*g*^ represents the expression profile of spot *i*. A spatial graph *G*_*s*_ = (*V, E*_*s*_) is constructed from the given spot coordinates **C** to define local spatial neighborhoods, where the vertex set *V* = {*v*_1_, … , *v*_*n*_} indicates tissue spots, and the edges *E*_*s*_ connect spots within a distance. The adjacency **A**_*s*_ of the spatial graph *G*_*s*_ is defined as *a*_*ij*_ = 1*/*(*d*_*ij*_)^*τ*^ if *d*_*ij*_ ≤ *r* and *i* ≠ *j*, and *a*_*ij*_ = 0 otherwise, where *d*_*ij*_ is the Euclidean distance between spots, *r* is the hyper-parameter of spatial radius, and *τ >* 0 controls the distance decay.

Given gene expression **G** and histological image ℐ, modality-specific affinity graphs *G*_*p*_ and *G*_*h*_ are constructed based on pathway activity and histological features, respectively. The pathway affinity graph *G*_*p*_ = (*V* , *E*_*p*_) captures pathway-level similarity relationships between spots. For each spot *i*, pathway activity vector **p**_*i*_ = R_*f*_ (**g**_*i*_) is computed by a pathway scoring function R_*f*_ : ℝ^*g*^ → ℝ^*p*^ (e.g., PaaSC [8]) that maps *g* genes to *p* functional pathways. An edge *e*_*ij*_ ∈ *E*_*p*_ encodes the pathway-level similarity between spots *i* and *j*. This relationship is quantified by the adjacency matrix **A**_*p*_ ∈ ℝ^*n×n*^, where each entry *A*_*ij*_ represents the pairwise similarity between **p**_*i*_ and **p**_*j*_. The graph structure is characterized by the normalized Laplacian 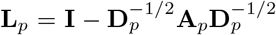, where **D**_*p*_ denotes the degree matrix. The pathway spectral embedding 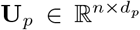 provides a low-dimensional representation of functional relationships, consisting of the *d*_*p*_ smallest non-trivial eigenvectors of **L**_*p*_.

The histology affinity graph *G*_*h*_ = (*V, E*_*h*_) captures morphological similarity between spots. For each spot *v*_*i*_, an image patch of size *s* × *s* centered at (*x*_*i*_, *y*_*i*_) is cropped from *I* and encoded into a histology feature vector using a pretrained encoder (e.g., UNI [4]). The adjacency matrix **A**_*h*_ ∈ ℝ^*n×n*^ quantifies edge similarities *E*_*h*_, where each entry *A*_*ij*_ represents the pairwise similarity between histology feature vectors of spots *i* and *j*. From **A**_*h*_, the normalized Laplacian **L**_*h*_ and the corresponding spectral embedding 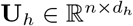 are computed following the same procedure as for the pathway graph.

Then, spatial diffusion captures higher-order connectivity across the tissue by propagating the spatial context. Specifically, a Personalized PageRank diffusion operator aggregates multi-hop spatial neighborhoods with decaying weights:

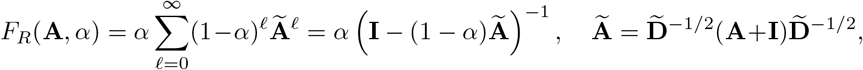

where 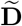 is the degree matrix of (**A** + **I**), and *α* ∈ (0, 1) controls the diffusion rate. The element values of the output matrix of *F*_*R*_(**A**_*s*_, *α*), which are smaller than a threshold *ϵ*, are set to zeros to maintain sparsity.

Given the pathway and histology spectral embeddings **U**_*p*_ and **U**_*h*_, spatially refined representations are obtained as:

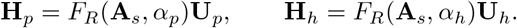

The spatially refined embeddings from pathway and image modalities are concatenated to form a combined representation, 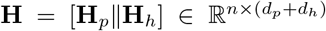, where [·∥·] denotes concatenation. Then, a joint cross-modal affinity graph *G*_*c*_ = (*V, E*_*c*_) is constructed based on the combined representation **H**. The affinity matrix **A**_*c*_ ∈ ℝ ^*n×n*^ quantifies the pairwise similarity between the integrated feature vectors of two spots *v*_*i*_ and *v*_*j*_. This graph encodes cross-modal similarities that integrate both pathway and morphological information. The diffusion operator refines the integrated representation, effectively combining functional and morphological relationships:

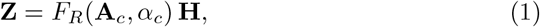

where *α*_*c*_ denotes the diffusion parameter for cross-modal integration. Finally, *k*-means clustering is applied to the integrated embedding **Z** to identify biologically meaningful spatial domains.

## 3 Experimental Results

We assessed our method on four gold-standard benchmark datasets generated by the 10x Genomics Visium platform, each providing spatially resolved gene expression profiles and H&E-stained histology images. The benchmark datasets include two human breast cancer datasets (BAS1 and DCIS), the mouse brain anterior data set (MBAD), and human dorsolateral prefrontal cortex (DLPFC). The two breast cancer datasets, BAS1 (Block A, Section 1) and DCIS (ductal carcinoma in situ), represent complex tumor microenvironments in fresh-frozen and formalin-fixed paraffin-embedded tissues, respectively. The MBAD set represents a coronal section of the adult mouse brain, which captures neuroanatomical regions. The human DLPFC profiles the layered prefrontal cortex, and we used sample 151673, which is a standard reference in spatial transcriptomics studies. We considered the state-of-the-art models, covering image-assisted representation learning (DeepST [15]), graph autoencoder-based embedding (STA-GATE [5]), contrastive learning (GraphST [10]), and pathway-informed multi-view graph modeling (PathMGCN [18]). Hyperparameters for all the benchmark methods, including neighborhood size and graph construction radius, were tuned based on the optimal settings recommended on the corresponding datasets in their original publications. Otherwise, we empirically optimized the hyperparameters. For our model, we empirically optimized hyperparameters including the diffusion transport rate and image patch size. We obtained curated pathways from MSigDB to computed pathway scores. We considered species-specific pathway sets corresponding to human and mouse of the datasets. For the clustering numbers, we set the numbers (*k*) of spatial domains as the ground truth numbers annotated by pathologists, when ground truths were available. We set *k*=20 for BAS1 [16], *k*=52 for the MBAD, and *k* = 7 for the DLPFC. For DCIS where ground truths were not available, we set *k*=10, which is the most frequently used setting. Note that most studies using DCIS have considered 10 ≤ *k* ≤ 15.

To compare performance, we computed various evaluation metrics, including Adjusted Rand Index (ARI), Silhouette Coefficient (SC), Davies-Bouldin index (DB), and Calinski-Harabasz index (CH) for the clustering task. ARI measures agreement between predicted clusters and reference annotation, when ground truths are available. SC evaluates intra-cluster cohesion versus inter-cluster separation, and DB measures cluster compactness and separation. CH compares between-cluster to within-cluster dispersion. Higher ARI, SC, and CH scores, and lower DB scores, indicate improved clustering results. We repeated the experiment ten times for reproducibility.

### 3.1 Performance Comparison with Benchmark Methods

BIGraph-ST demonstrated robust performance and notable improvements on most benchmark datasets (Table 1). BIGraph-ST achieved significantly superior clustering accuracy for cancer tissues. For the human breast cancer BAS1, BIGraph-ST outperformed all competing methods with an ARI of 0.71 ± 0.01, representing a 24.5% improvement over the second-best (0.57; *p <* 0.01, Wilcoxon signed-rank test). Similarly, BIGraph-ST ranked first across all metrics on the human breast cancer DCIS (SC: 0.37, DB: 1.04, CH: 938), improving the SC by 12.1% compared to the runner-up. For MBAD, BIGraph-ST achieved a competitive ARI of 0.45 ± 0.01 (4.7% higher than the second-best with 0.43) and ranked second in SC, DB, and CH following PathMGCN. On DLPFC, GraphST achieved the highest ARI (0.60 ± 0.01) and BIGraph-ST achieved the (0.53 ± 0.01). The superior performance on cancer datasets (BAS1 and DCIS) may suggest that BIGraph-ST effectively captures disease-associated spatial heterogeneity at the functional level, whereas the relatively lower performance on DLPFC may be attributable to its continuous cortical layering rather than discrete boundaries.

**Table 1:**
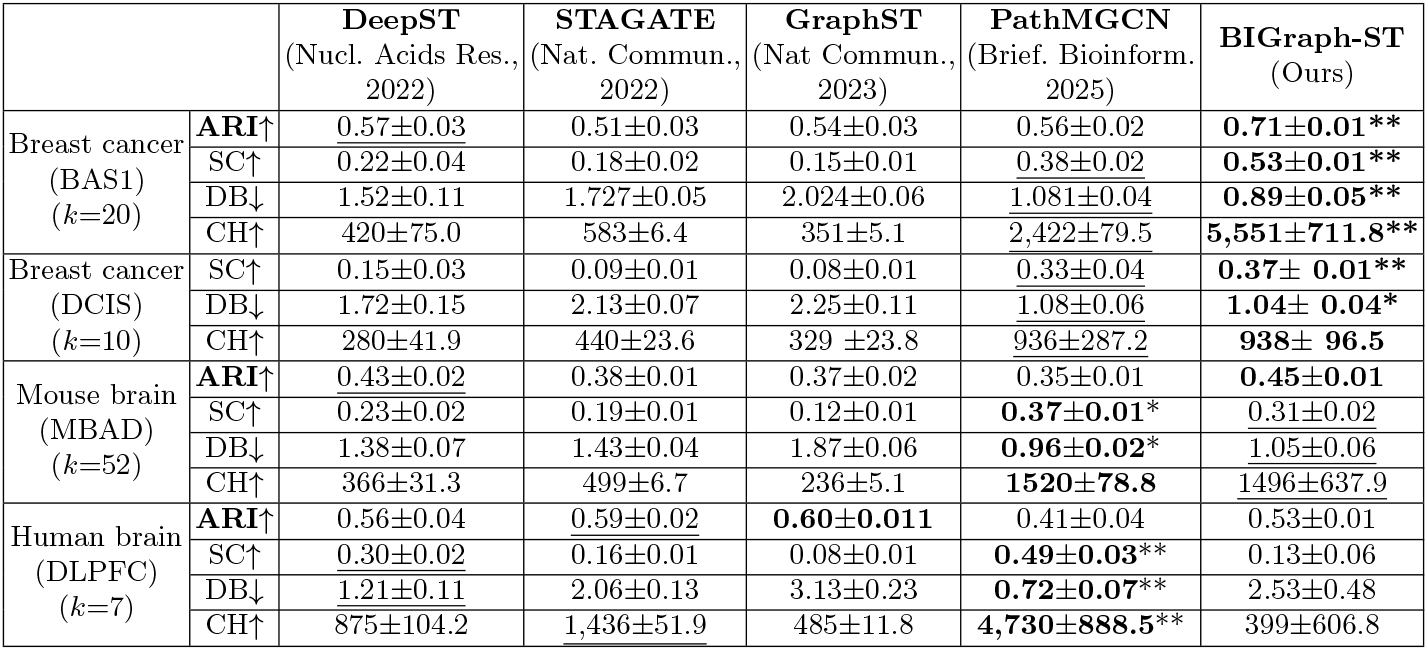
Performance comparison with state-of-the-art methods. The best performance is highlighted in **bold**, and the second-best is indicated in <u>underline</u> with statistical significance (Wilcoxon signed-rank: ^*^ (*p <* 0.05) , ^**^ (*p <* 0.01)).

### 3.2 Ablation Study

We investigated the contribution of each component of BIGraph-ST by selectively removing or replacing the key modules, including pathway representation and histological information (Table 2). Compared to the baseline (Gene + Image), incorporating pathway scores (PaaSC + Gene + Image) yielded ARI improvements of 11.4% on MBAD (0.440 vs. 0.395), 10.4% on BAS1 (0.710 vs. 0.643), and 35.8% on DLPFC (0.524 vs. 0.386). Relative to the gene-only baseline, incorporating histological image features increased ARI by 13.5% on MBAD (0.395 vs. 0.348) and 14.4% on BAS1 (0.643 vs. 0.562), and 11.2% on DLPFC (0.386 vs. 0.347). We further compared two pathway scoring strategies, PaaSC [8] and GSVA [6]. PaaSC consistently outperformed GSVA under the same multi-modal setting, yielding relative ARI gains of 7.3%, 5.5%, and 2.7% on the MBAD, BAS1, and DLPFC datasets, respectively.

**Table 2:**
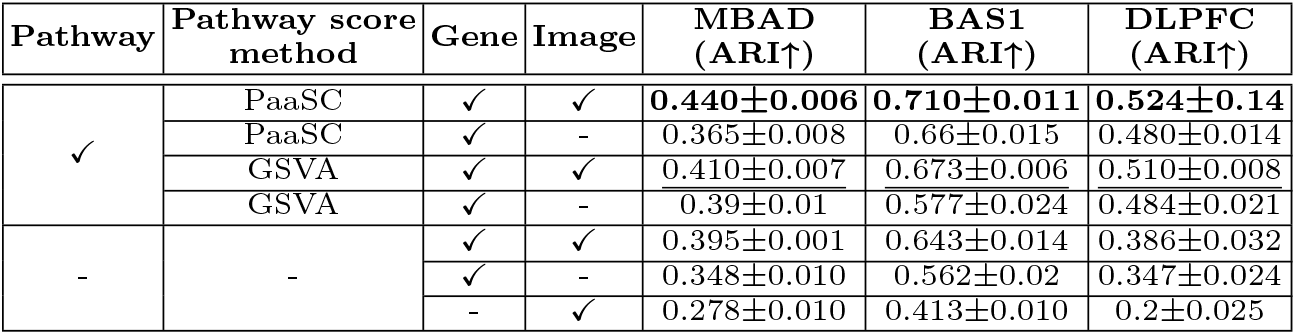
Ablation experimental results. The best performance is highlighted in bold, and the second-best is indicated in <u>underline</u>.

| Pathway | Pathway score method | Gene | Image | MBAD (ARI↑) | BAS1 (ARI↑) | DLPFC (ARI↑) |
| --- | --- | --- | --- | --- | --- | --- |
| ✓ | PaaSC | ✓ | ✓ | **0.440±0.006** | **0.710±0.011** | **0.524±0.14** |
| | PaaSC | ✓ | - | 0.365±0.008 | 0.66±0.015 | 0.480±0.014 |
| | GSVA | ✓ | ✓ | 0.410±0.007 | 0.673±0.006 | 0.510±0.008 |
| | GSVA | ✓ | - | 0.39±0.01 | 0.577±0.024 | 0.484±0.021 |
| - | - | ✓ | ✓ | 0.395±0.001 | 0.643±0.014 | 0.386±0.032 |
| | | ✓ | - | 0.348±0.010 | 0.562±0.02 | 0.347±0.024 |
| | | - | ✓ | 0.278±0.010 | 0.413±0.010 | 0.2±0.025 |

### 3.3 Model Interpretation

We analyzed the functional domains identified by BIGraph-ST using BAS1. BIGraph-ST partitioned BAS1 into 20 spatial domains that closely correspond to pathologist-annotated invasive ductal carcinoma (IDC), ductal/lobular carcinoma in situ (DCIS/LCIS), tumor edge, and healthy regions (Fig. 2). We identified signature pathways for each spatial domain by performing a Student’s *t*-test to compare pathway activity scores between the target domain and the rest, and ranking pathways according to statistical significance (Table 3). The resulting domain-signature pathways revealed coherent biological domains. For example, domain 4 corresponded to the healthy region and these signature pathways are indicative of a benign stromal compartment. Domain 9 corresponded to the DCIS/LCIS compartment, where signature pathways highlight extracellularmatrix and glycosphingolipid remodeling within a pre-invasive epithelial context. Domain 6 represented the IDC tumor core, marked by broader metabolic and signaling programs consistent with the high metabolic demands of invasive malignancy. BIGraph-ST refinesd tumor edge annotations into distinct microenvironmental bands (e.g., domains 10, 15, 17, 19). For instance, domain 10 showed tumor-associated macrophage signaling, whereas domain 15 was dominated by CD4 T cell activity [1]. These results indicate that BIGraph-ST captures both cell-type–associated signals and key cellular processes at the domain level, enabling biologically interpretable spatial characterization.

**Table 3:** Signature pathways for representative spatial domains in BAS1.

|  | Domain 4 | Domain 9 | Domain 6 | Domain 10 | Domain 15 |
| --- | --- | --- | --- | --- | --- |
| **Matched annotation** | Healthy_1 | DCIS/LCIS_4 | IDC_2 | Tumor edge_6 | Tumor edge_3 |
| **Top 3 pathways** | Regulates expression of components of tight junctions | Glycosaminoglycan degradation | Amino acid conjugation | Platelet pathway | Binding and uptake of ligands by scavenger receptors |
|  | Regulation of cytoskeletal remodeling | Glycosphingolipid metabolism | Iron metabolism disorders | Creation of C4 / C2 activators | GCR pathway |
|  | Rho pathway | Glycosphingolipid catabolism | Microtubule RhoA signaling pathway | Trafficking and processing of endosomal TLR | CBL pathway |
| **Summary** | Non-malignant stroma | ECM / glycosphingolipid remodeling | Metabolic and signaling programs | Tumor-associated-macrophages | Immune cell signaling |

**Fig. 2:**
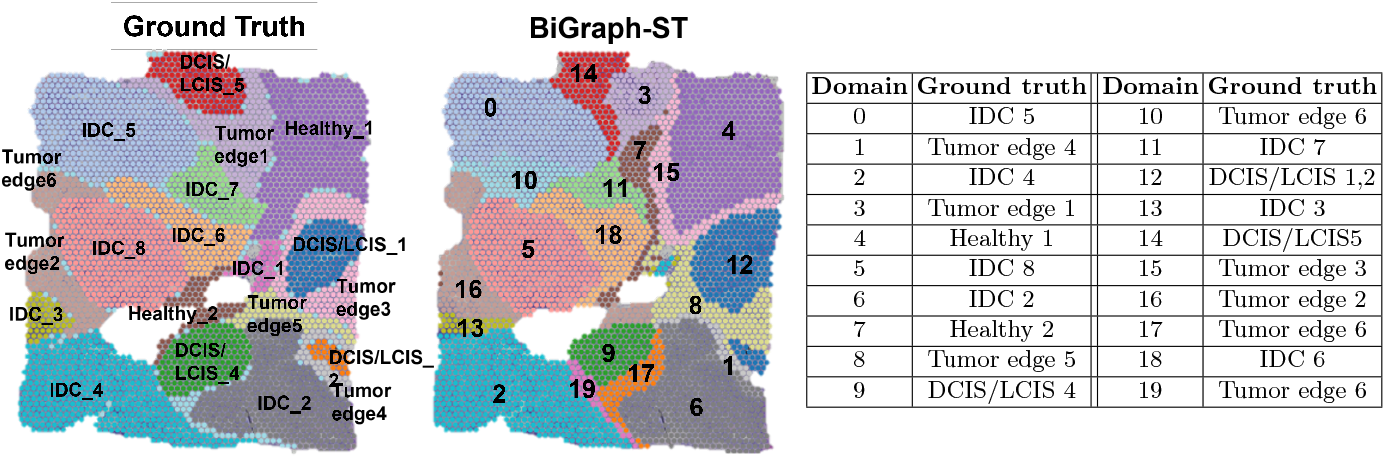
BIGraph-ST domains and their corresponding ground-truth on BAS1.

## 4 Conclusion

In this study, we introduce a novel multimodal framework for spatial domain identification that integrates pathway-level transcriptomic information and histological image features, while explicitly modeling high-order spatial connectivity. The experimental results demonstrated that BIGraph-ST achieved superior performance compared with state-of-the-art methods on most datasets, and ablation studies further confirmed the contribution of pathway level representation and multi modality integration. Furthermore, biological interpretation on the human breast cancer reveals domain-specific functional patterns, supporting the biological relevance of the identified spatial domains. Overall, our study provides a robust and interpretable approach for identifying biologically meaningful spatial domains and offers a valuable tool for gaining biological insights into complex tissue microenvironments.

## Notes

### Competing Interest Statement

The authors have declared no competing interest.

